# Meta-analysis of livestock effects on tree regeneration in oak agroforestry systems

**DOI:** 10.1101/2024.03.02.583097

**Authors:** Abdullah Ibne Wadud, Miguel N. Bugalho, Pedro Gonçalves Vaz

**Author notes:** **Correspondence:** Pedro G. Vaz.

## Abstract

Livestock grazing occupies over a quarter of terrestrial land and is prevalent to agroforestry ecosystems, potentially affecting the survival, growth, and density of trees’ early developmental stages, such as seeds, seedlings, and saplings. To address the effects of livestock on tree recruitment in the face of ongoing debates about their impacts, we conducted a 33-year meta-analysis in *Quercus*-dominated agroforestry systems. Our analysis revealed a consistently negative effect of livestock on oak acorns, seedlings, and saplings. Significantly, livestock body size influenced oak regeneration, with small-sized livestock, notably sheep and goats, having a more pronounced negative impact compared to mixed-size systems, mainly involving cattle and sheep. The effects of small-sized livestock were markedly detrimental on acorn survival and seedling/sapling density, although no studies eligible for meta-analysis examined large livestock impacts on acorns. Overall, mixed-size livestock systems, often involving cattle and sheep, lessen the negative effects. Our findings indicate that the body size and foraging behaviors of livestock should be considered for the ecological sustainability of the tree component in agroforestry systems. While protective measures have long been integral to well-managed agroforestry systems, our results underscore the importance of integrating diverse livestock sizes and applying specific protective strategies, particularly for acorns and saplings, to further refine these practices. Future research should expand to underrepresented regions and livestock types to refine global agroforestry management practices.

**Highlights:**

- Livestock reduces oak tree regeneration at acorn, seedling, and sapling stages.
- Small livestock effect more negative, especially acorn survival and young oak density.
- Mixed-size livestock agroforestry systems reduce harm more than small single-size ones.
- We challenge perceptions that replacing cattle with sheep is necessarily less harmful.
- We advocate for diverse livestock sizes in *Quercus*-dominated agroforestry systems.

## 1. Introduction

Livestock grazing occurs on over a quarter of terrestrial land (Taylor and Rising, 2021; FAO, 2023) and is common in agroforestry ecosystems, which merge forestry with agriculture and frequently also with livestock production (Allen et al., 2011). In these ecosystems, livestock of different body sizes and foraging behaviors affect key tree reproductive stages such as seed, seedling, and sapling, altering survival, growth, and density of new tree recruits (Pulido and Díaz, 2005). Thus, managing livestock in agroforestry systems requires balancing production with potential negative effects of livestock on tree life stages (Brown et al., 2018). Analyzing the effects of livestock on tree survival across similar agroforestry systems worldwide may provide better informed sustainable management strategies.

*Quercus*-dominated agroforestry and silvopastoral systems occur in various parts of Europe, including the Iberian Peninsula (e.g., dehesas and montados), east-central and north-central Europe, as well as in North Africa, North America (e.g., California and Texas), and in Asia (Plieninger et al., 2015; Tantray et al., 2017). These systems span semi-natural and managed landscapes across temperate regions and are particularly valued for their socio-economic and cultural significance as well as associated biodiversity (Pantera et al., 2018; Stavi et al., 2022). They provide a plethora of ecosystem goods including fuelwood, game, crops, and cork (Bugalho et al., 2009), alongside services like biodiversity conservation (Plieninger et al., 2011). Regrettably, they face threats from climate change (Príncipe et al., 2019; Díaz et al., 2021), pathogens (Branco and Ramos, 2009; Brasier, 1992, 1996), phytophagous insects (Branco et al., 2002), wildfires (Moreira et al., 2011; Vaz et al., 2013), and overgrazing (Vaz et al., 2019). These threats contribute to low oak recruitment widely documented across Europe (Pulido et al., 2001), North America (Rogers et al., 1993), North Africa (Campos et al., 2007), and the Middle East (Dufour-Dror, 2007). Particularly, the impacts of livestock grazing on various early oak life stages remain unsubstantiated globally. Regarding tree regeneration, the process of recruiting individuals to sustain the adult population and counteract mortality losses (Harper, 1977), the impact of livestock on early life stages varies from expected negative (López-Sánchez et al., 2014) to positive effects (Leiva et al., 2022) or even mixed outcomes (Laskurain et al., 2013), with livestock body size and foraging behaviors being important factors (e.g., Ball and Tzanopoulos, 2020). Moreover, while the detrimental effects of overgrazing are well documented, management neglect and grazing abandonment also pose significant threats to the sustainability of oak silvopastoral systems (Bugalho et al., 2011; Plieninger et al., 2015). Overall, the effects of livestock on oak regeneration, including both their signal and magnitude, continue to stir ongoing debate.

Variability in livestock effects on acorn, seedling, and sapling stages, along with differing metrics such as survival, growth, and density, drive the ongoing debate. Acorn survival (viable acorns to become seedlings) is limited by livestock consumption and soil compaction. Yet, livestock activity can bury and aid acorn germination (Leiva and Sobrino-Mengual, 2022). Livestock hinder animal-mediated acorn dispersal (Vaz et al., 2024) but also consume those infested by pests (Canelo et al., 2021). On seedlings, the impact is also complex and fuels the debate. Trampling leads to soil compaction, bare soil, altered litter quality, enhanced soil erosion, reduced nitrogen-fixing species, and hindered seedling establishment (Vázquez, 2002; Etchebarne and Brazeiro, 2016; Cierjacks et al., 2004; Fortuny et al., 2020), reducing seedling density and opposing regeneration niches (Szewczyk and Szwagrzyk, 2010). Conversely, livestock reduce competition from annual grasses and may decrease insect herbivory (Tyler et al., 2008; Muñoz et al., 2009; Gallego et al., 2017), ultimately benefiting oak seedling survival (Zhang et al. 2019; Vaz et al. 2019). For saplings, frequent grazing and rubbing against them often delay growth and increase mortality (Roula et al., 2019), but moderate grazing deters shrub encroachment, mitigates wildfires (Rouet-Leduc et al., 2021), reduces competition with herbs and shrubs, and may facilitate growth in opening areas (Uytvanck et al., 2010; Reiner and Craig, 2011; Mazzini et al., 2018; Vaz et al., 2019). Notably, goats, often pointed out for the damage they cause to woody vegetation, can nevertheless be used to control invasive species and reduce fire hazards (Ruiz-Mirazo et al., 2011, Mena et al., 2016). Indeed, the maintenance of open habitats by livestock can play a critical positive role in oak recruitment (Bobiec et al., 2018; Wolański et al. 2021).

Beyond an overall effect of domestic livestock, the nuances of body size and foraging behaviors in agroforestry regeneration warrant further examination. In domesticated ruminants, a strong relationship between body size and foraging behavior is to be expected, despite exceptions such as cattle and sheep, which differ in size but are primarily grazers, preferentially feeding on grasses. However, importantly, within the same feeding strategy, differences in body size may lead to varied utilization of plant resources; for instance, sheep can graze closer to the ground than cattle, affecting the ecology of the system differently. On the other hand, smaller browsers like goats preferentially consume woody plants, such as oak seedlings and saplings (Laskurain et al., 2013; Abraham et al., 2018). Moreover, different livestock types can distinctly affect various life stages. Namely, small ruminants, particularly sheep and goats, may have levels of acorn consumption similar to those of wild ungulates (Leal et al., 2022). Particularly, goats are known for consuming acorns (Papachristou et al., 2005; Froutan et al., 2015) and are expected to be more harmful than grazing cattle and sheep, although these also consume acorns (Papachristou and Platis, 2011; Varga et al., 2020). Historically, goats have often been subject to regulations or even full exclusion because of their browsing behavior and its negative effects on trees and other woody vegetation (Humphrey et al., 1998; Vera, 2000). Considering the effects of livestock size on tree regeneration, some authors recommend smaller livestock like sheep over larger ones like cattle (e.g., López-Sánchez et al., 2016) within the same feeding strategy, as both are grazers. This, however, may not consider other effects such as acorn consumption. A comprehensive analysis of effect magnitudes across varied studies and conditions is essential to clarify whether livestock body size is a significant predictor of their impact on oak regeneration in agroforestry systems.

To address the variability in grazing effects, we conducted a meta-analysis of studies spanning over the last 33 years. This research synthesis compared grazed versus ungrazed areas in *Quercus*-dominated agroforestry systems to discern the influence of livestock on each of the early life stages of oak trees. We tested the following hypotheses. 1) Livestock have an overall negative combined effect on survival of acorns and survival, growth, and density of seedlings and saplings. 2) Livestock effects vary in sign and magnitude among acorn, seedling, and sapling stages. 3) The effect of livestock on early oak stages significantly vary with livestock size, with goats expected to cause more negative outcomes than mixed-size livestock systems. 4) The impact of grazing on oak life stages would differ according to the metric used, with expected less negative effects on survival and greater variability in growth and density metrics due to young oaks’ resilience, including their ability to resprout. With this meta-analysis, we not only elucidate the varied impacts of livestock on oak regeneration but also highlight areas for future research and offer insights for the effective conservation and management of *Quercus*-dominated agroforestry systems.

## 2. Materials and Methods

### 2.1. Literature review and inclusion criteria

We conducted a literature search using the Clarivate Analytics Web of Science^TM^ database, focusing on articles and review articles published in English between 1990 and July 2023. Our search string combined the keywords “(*Quercus* OR oak*) AND (regenerat* OR recruit* OR seedling* OR sapling* OR establish* OR acorn OR seed*) AND (graz* OR herd* OR pastur* OR brows* OR herbiv* OR predat*)”. To narrow down the search to relevant articles, we limited it to the research areas of Ecology, Forestry, Plant Sciences, Biodiversity Conservation, Environmental Sciences, Evolutionary Biology, Biology, and Agriculture Multidisciplinary. The search yielded 2,284 publications, which were subsequently refined following the steps outlined in the PRISMA protocol (Preferred Reporting Items for Systematic Reviews and Meta-analyses; Fig. 1). The refinement ensured the selection of studies eligible for meta-analysis (see Gerstner et al., 2017) that met the following inclusion criteria: (i) conducted in agroforestry systems dominated or subdominated by an oak tree species; (ii) addressing the effects of livestock; (iii) comprising both livestock grazed and ungrazed (control) treatments; (iv) investigating the effects on early tree life stages, including seeds, seedlings, or saplings; (v) and whose response variables included measurements of acorn survival, seedling and/or sapling survival, seedling growth, and seedling density.

**Fig. 1.**
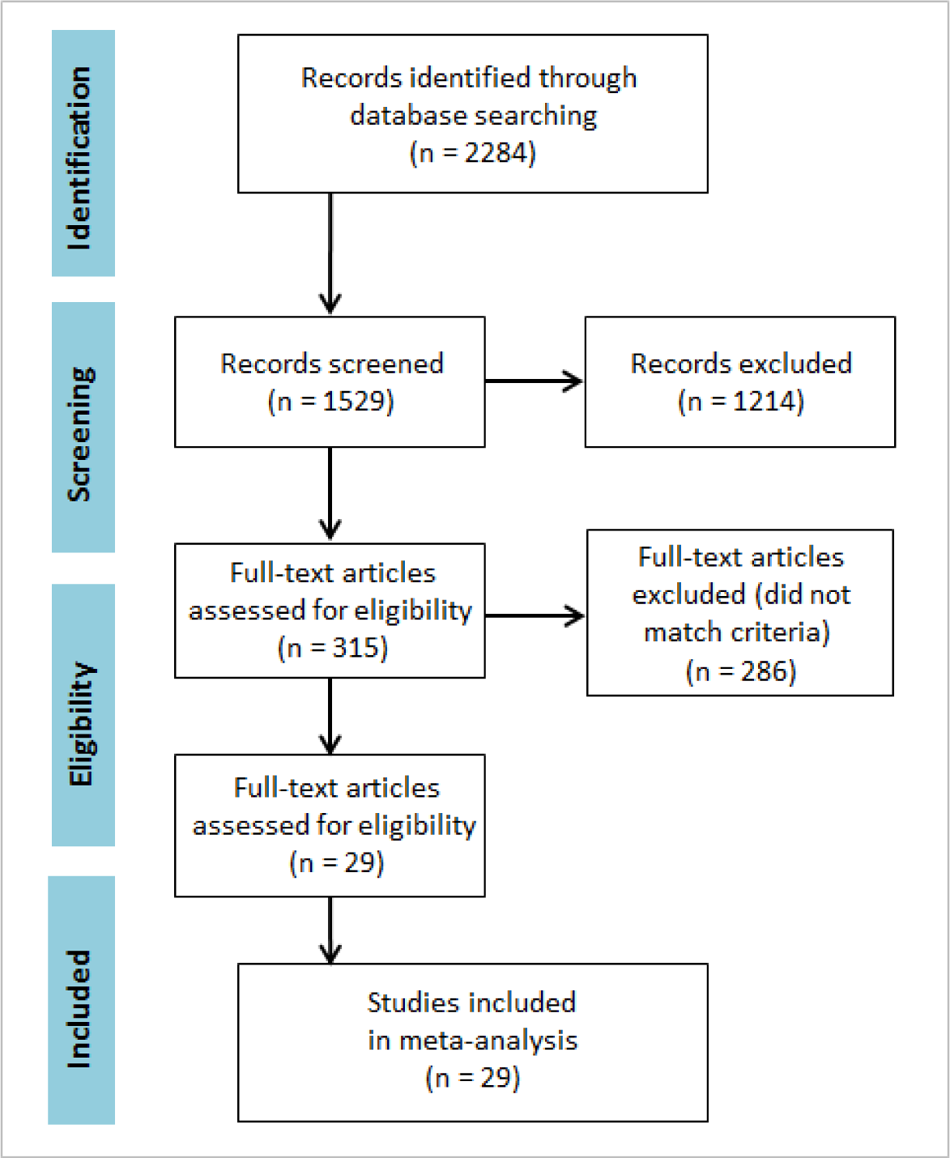
PRISMA diagram illustrating the study selection process.

To supplement the search, we thoroughly examined the reference lists of each retained article, which resulted in the inclusion of the following publications not captured by the initial search string criteria: Allred et al., 2012; Laskurain et al., 2013; and Pearse et al., 2014. In total, our search identified 54 case studies from 29 scientific articles that fulfilled the specified criteria.

### 2.2. Data extraction

We extracted the sample size, and then the mean and the standard deviation of the response variables per treatment from text, tables, or graphs. When not provided in the text or tables, we extracted values from graphs by zooming in on the screen. Survivals of seeds, seedlings, or saplings were expressed as percentages, calculated as (subjects at end ÷ initial subjects) × 100 if not directly provided. When not directly provided by the authors, we derived growth as the variation in height over time, and density as the number of individuals per square meter. We relied on the authors’ definitions to differentiate between seedlings and saplings of the same species, acknowledging minor variations in this classification across studies. When the included studies involved multiple levels of grazing treatment (e.g., Rossetti and Begella, 2014; Dorji et al., 2020), we calculated an average where applicable. To compare the effects of livestock in different oak agroforestry systems, we categorized each case study into two types: (1) oaks and (2) oaks with broadleaves and/or conifers. To compare livestock effects by animal size, we categorized the study cases as either large (cattle, horses), small (sheep, goats), or mixed. Thus, a case study in our meta-analysis was defined as a unique combination of agroforestry type (oaks, oaks with broadleaves and/or conifers), livestock size (small, mixed, large), life stage (acorn, seedling, sapling), and response variable (survival, growth, density).

Among 29 scientific articles (Table 1), 12 featured one case study, 12 contained two, and 2 included three. Three articles (Rossetti and Begella, 2014; Costa et al., 2017; and Murphy et al., 2021) each contributed four case studies, which raises potential issues of pseudo-replication across a total of 54 case studies (Massad and Dyer, 2010). In some instances, we combined results from different time periods (years, months) presented separately in the study, if the data were collected in the same study areas.

**Table 1.** Summary of articles. Livestock: S = sheep; G = goat; C = cattle; H = horse. Oak life stage: Se = seedling; Sa = sapling; Ac = acorn. Response: De = density; Gr = Growth; Su = survival. Tree species: *Q*. = *Quercus*; other = other broadleaves and/or conifers.

| Study | Country | No. cases | Livestock | Stage | Response | Tree species |
| --- | --- | --- | --- | --- | --- | --- |
| Reiner & Craig 2011 | USA | 1 | S, C | Se | De | *Q. douglasii* |
| Clements et al. 2011 | Canada | 1 | S | Se | Gr | *Q. garryana* |
| Ruiz-Mirazo & Robles 2012 | Spain | 2 | G | Se, Sa | Gr | *Q. ilex* |
| Espelta et al. 2006 | Spain | 2 | C | Sa | Gr | *Q. ilex*, *Q. cerrioides* |
| Abraham et al. 2018 | Greece | 1 | S, G | Se | De | *Q. frainetto* |
| Plieninger et al. 2004 | Spain | 2 | S, G | Se | De | *Q. ilex* |
| DufourDror et al. 2007 | Israel | 1 | S, G, C | Se | De | *Q. ithaburensis* |
| Rossetti & Bagella et al. 2012 | Italy | 4 | S | Se | Gr, De | *Q. suber* |
| Leiva & FernandezAles 2003 | Spain | 2 | S | Ac | Su | *Q. ilex* |
| Pulido et al. 2010 | Spain | 1 | S | Ac | Su | *Q. ilex* |
| Taylor et al. 2008 | USA | 2 | S, C | Ac | Su | *Q. lobata*, *Q. agrifolia* |
| Arosa et al. 2015 | Portugal | 2 | S, C, S | Ac | Su | *Q. ilex*, *Q. suber* |
| Davis et al. 2011 | USA | 1 | S, C | Ac | Su | *Q. lobata* |
| Cierjacks & Hensen 2004 | USA | 1 | S, G | Ac | Su | *Q. ilex* |
| Leiva & Vera 2015 | Spain | 3 | S, G, H | Ac | Su | *Q. ilex*, *Q. suber* |
| Allred et al. 2012 | USA | 2 | S, G, C | Se | De | *Q. fusiformis* |
| Pearse et al. 2014 | USA | 3 | S, C | Se | De | *Q. lobata* |
| Leiva & Sobrino-Mengual 2022 | Spain | 2 | C | Se | Su | *Q. ilex* |
| Diaz-Hernandez et al. 2021 | Spain | 2 | S, C | Ac | Su | *Q. ilex* |
| Parsons et al. 2021 | USA | 1 | C | Se | De | *Q. lobata*, *Q. douglasii*, *Q. kelloggii* |
| Dorji et al. 2020 | Bhutan | 2 | C | Se | Su, Gr | *Q. semecarpifolia* |
| Costa et al. 2017 | Spain | 4 | S, C | Se | Su | *Q. pyrenaica*, *Q. ilex* |
| Murphy et al. 2021 | England | 4 | S, G | Se, Sa | Su, Gr, De | *Q. robur*, *Q. petraea* |
| Roula et al. 2019 | Algeria | 1 | S, C | Sa | Gr | *Q. suber* |
| Phillips et al. 2007 | USA | 1 | S, C | Se | Gr | *Q. douglasii* |
| Laskurain et al. 2013 | Spain | 1 | S | Se | Su | *Q. robur*, other |
| Moradi et al. 2021 | Iran | 1 | S, G | Sa | Gr | *Q. brantii*, other |
| Fortuny et al. 2020 | France | 2 | C | Se | De | *Q. pubescens*, other |
| Amsten et al. 2021 | Sweden | 2 | S, G, H | Sa | Su, Gr | *Q. robur*, other |

### 2.3. Analyses

For each of the 54 study cases, we used Hedges’ *d* metric and a confidence interval (CI) to estimate effect sizes (Hedges, 1981; Gurevitch and Hedges, 2001). The analyses were performed using the *metafor* package (version 3.8-1; Viechtbauer, 2010) in R (version 4.2.2; R Core Team, 2020). Hedges’ *d* represents the standardized mean difference between areas with and without livestock. Negative values indicate lower survival for acorns, seedlings, and saplings, as well as reduced seedling growth and sapling density in grazed areas compared to control areas. Positive values indicate the opposite. We considered an effect size as significant if its 95% CI did not overlap with zero (Koricheva et al., 2013). Effect sizes are commonly interpreted using Cohen (1988)’s criteria: <|0.2| is low, |0.2–0.5| is moderate, >|0.8| is high, and >1.0 is very high. To account for potential pseudo-replication and dependencies among case studies reported within the same study, we used the *robust* function from the *metafor* package, which provides robust variance estimates when multiple effect sizes are derived from the same article (Hedges et al., 2010).

We used random-effects meta-analyses to estimate mean effect sizes across pools of case studies. This approach assumes that the effect sizes in case studies are a random sample of all possible effect sizes (Borenstein et al., 2010). First, we estimated the grand mean effect size and 95% CI across all studies to see if livestock had a general effect on all the response variables combined (Koricheva et al., 2013). Then, to measure the consistency across studies, we calculated the among-studies heterogeneity (τ² and associated *Q* statistics). To account for the dependence of τ² on sample size, we also calculated *I*^2^, a standardized estimate of total heterogeneity ranging from 0 to 1 (Nakagawa et al., 2017; Borenstein, 2022). To assess whether there was evidence of an overall effect of livestock on a particular group of response variables (e.g., survival, combining all the early tree life stages), we calculated omnibus tests (e.g., Moreira et al., 2019). Last, we assessed publication bias (Appendix A) using funnel plots (Fig. A.1, Appendix A) and Rosenthal’s fail-safe numbers (Koricheva et al., 2013). Publication bias is a critical concern in meta-analysis, as studies with statistically significant results are often more likely to be published than those with nonsignificant findings (Fragkos et al., 2014).

## 3. Results

### 3.1. Overview of the case studies

Most case studies eligible for meta-analysis were conducted in Europe (n = 37), followed by the Americas (12), Asia (4), and Africa (1) (Fig. B.1, Appendix B). Eleven of the 54 case studies were carried out in the USA. Spain recorded the highest number of cases in Europe (22). The 54 case studies were extracted from peer-reviewed articles published in 22 JCR-listed journals. Among them, 10 ranked in Q1, 12 in Q2, and 5 in Q3 quartile within JCR subject categories. Nearly half of the case studies (44%) assessed the effects of mixed-size livestock, often including cattle and sheep (67% of mixed-size studies) or a combination of cattle, sheep, and goats (12%), with some featuring horses, sheep, and goats (21%). In the 39% of small-sized livestock studies, 52% examined only sheep, 9% only goats, and 38% both sheep and goats. All the large-sized livestock case studies (17%) involved cattle. Seedlings were the most common oak life stage in the case studies (48%), followed by acorns and saplings (26% each).

Forty-four percent of the studies measured livestock effects on the survival of early life stages (acorn, seedling, or sapling), while 31% and 24% assessed density and growth in seedlings and saplings, respectively. Forty-eight of the 54 case studies focused on oak agroforestry systems and six included oaks with broadleaves and/or conifers.

### 3.2. Combined survival, growth, and density effects

Livestock had a combined negative impact on the early life stages of oaks in agroforestry systems, as measured by survival of acorns, seedlings, and saplings, and density and growth of seedlings and saplings (Fig. 2; Fig. B.2, Appendix B). The combined effect size, as represented by Hedge’s *d*, was −0.87 (95% CI: [−1.12, −0.62]), indicating a high negative effect. The effect was negative, regardless of continent or country. The global meta-analysis showed a substantial amount of total heterogeneity (τ² = 0.73, *Q*T = 4260.8, *P* < 0.001), 98% of which was attributable to between-study heterogeneity (*I^2^* = 98%).

**Fig. 2.**
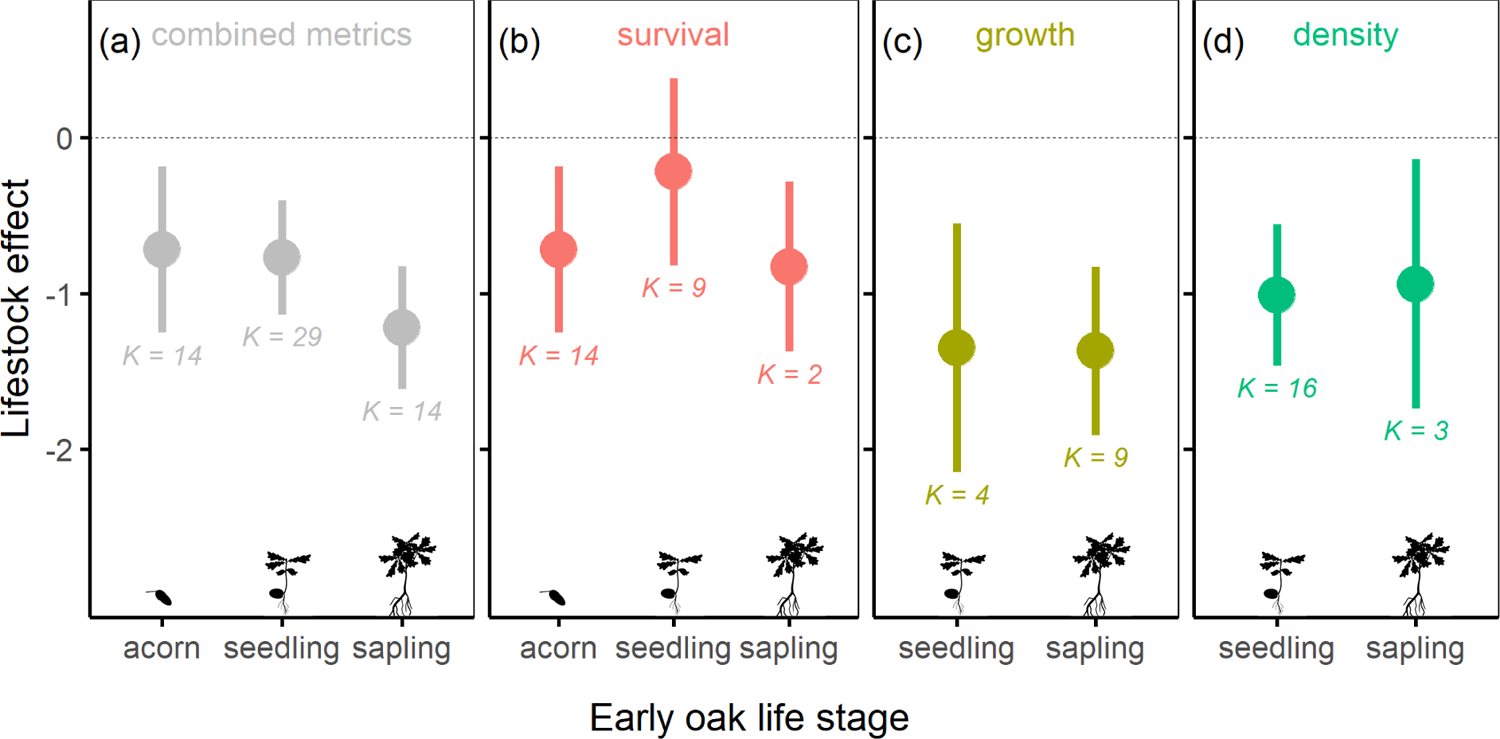
Mean effect size (Hedges’ *d*) of domestic livestock on combined survival, growth, and density effects across life stages (a), and on survival, growth, and density individually (b, c, d). Dots with error bars represent model parameter estimates and their 95% confidence intervals (CI). ‘*K*’ denotes the number of case studies. The mean effect size is considered significant if the 95% CI does not intersect the dashed zero line (no effect).

#### 3.2.1. Effects by oak early life stage

Livestock had a consistently negative impact on oak regeneration across all stages: acorn (*Q* = 3008, *P* < 0.001), seedling (*Q* = 404, *P* < 0.001), and sapling (*Q* = 505, *P* < 0.001). The effect was not deemed significantly different among these life stages (*Q*M = 2.8, *P* = 0.249) but it uniquely reached ‘very high’ severity on saplings (Cohen, 1988). Variability between studies was relatively high in all three stages (τ² > 0.43 in each case), with over 90% of this variability attributed to differences between case studies within each stage.

#### 3.2.2. Effects by livestock size

Livestock effects varied significantly by size (*Q*M = 11.92, P = 0.001), but were negative across categories (Fig. 3a; Fig. B.3, Appendix B). The most pronounced negative effect was observed with small-sized livestock (−1.37; [−1.81, −0.92]). In comparison, large-sized and mixed-sized livestock systems tended to exhibit less negative effects (*Z* = 1.8, *P* < 0.072 and *Z* = 3.4, *P* < 0.001, respectively). The mixed-size livestock subgroup showed lower heterogeneity in effect sizes (τ² = 0.26) and more than 97% of the variability was due to differences between case studies in all size subgroups.

**Fig. 3.**
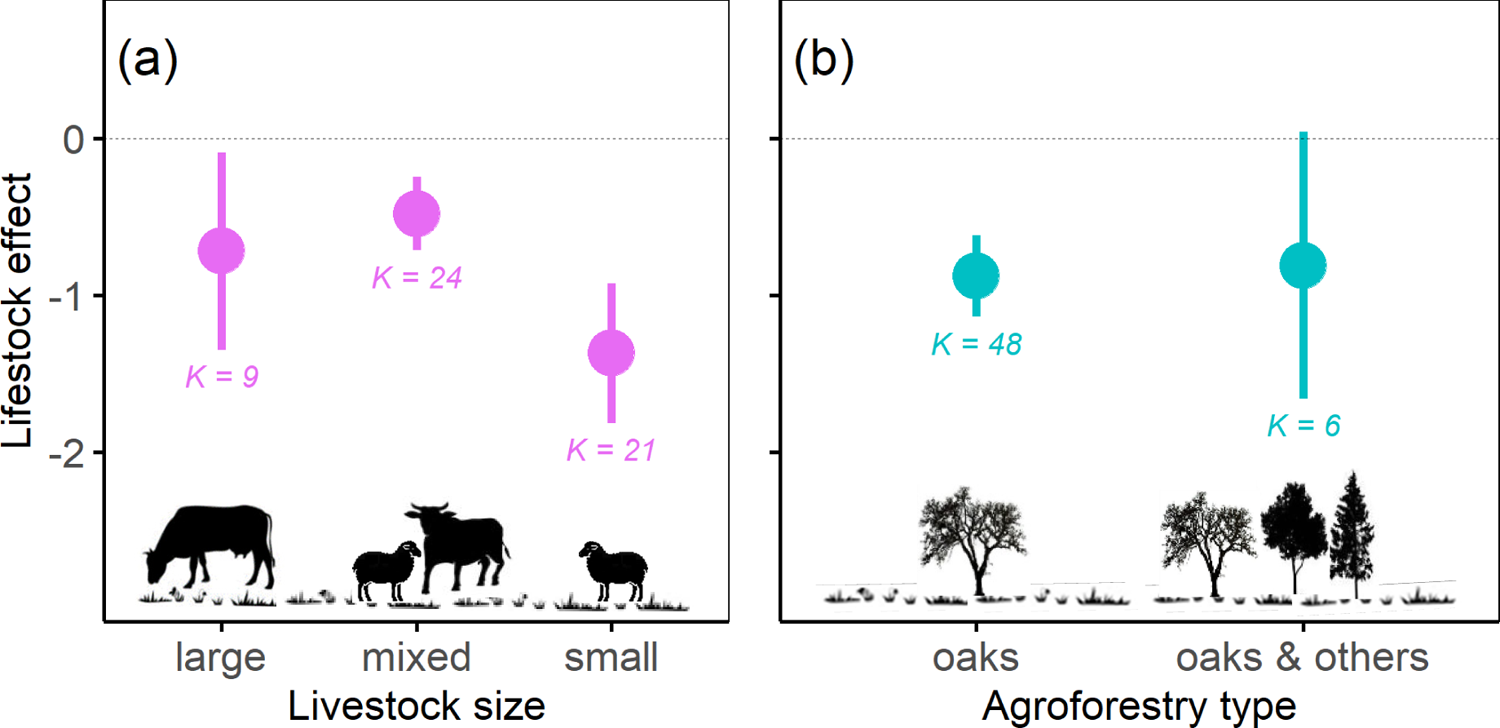
Mean effect size of domestic livestock on the early life stages of oaks, categorized by body size (a) and type of agroforestry system (b; either oak trees only or oaks mixed with broadleaves and/or conifers). Refer to Figure 2 for more explanations.

#### 3.2.3. Effects by oak agroforestry type

The effects of livestock did not vary between case studies carried out in agroforestry systems with oaks only and systems with oaks and broadleaves and/or conifers (*Q*M = 0.03, *P* = 0.857) (Fig. 3b). The effects were significantly negative in both the first (*Q* = 33, *P* < 0.001) and second types of system (*Q* = 87, *P* < 0.001). In both types, substantial heterogeneity in effect sizes was observed (τ² > 0.71), with over 98% of the variability in effect sizes within each type attributable to between-study differences.

### 3.3. Effects by type of response to livestock

#### 3.3.1. Effects on early oak survival

Of the 25 case studies examining the impact of livestock on early oak survival in agroforestry systems, an overall negative effect was found (−0.57; [−0.94, −0.19]), without dependence on the life stage (*Q*M = 1.6, *P* = 0.459). However, the negative effect was less pronounced at the seedling stage (−0.22; [−0.82, 0.38]) and was observed in the only two case studies concerning the oak sapling stage (−0.83; [−1.37, −0.28]). The 14 case studies focusing on acorn survival provided clear evidence of a negative effect (−0.72; [−1.25, −0.18]), with 5 involving small-sized and 9 involving mixed-size livestock. Further analysis demonstrated that the impact on acorn survival was significantly more negative in case studies with small-sized livestock compared to those with mixed-size livestock (*Q*M = 11.5, *P* < 0.001) (Fig. 4; Fig. B.4, Appendix B).

**Fig. 4.**
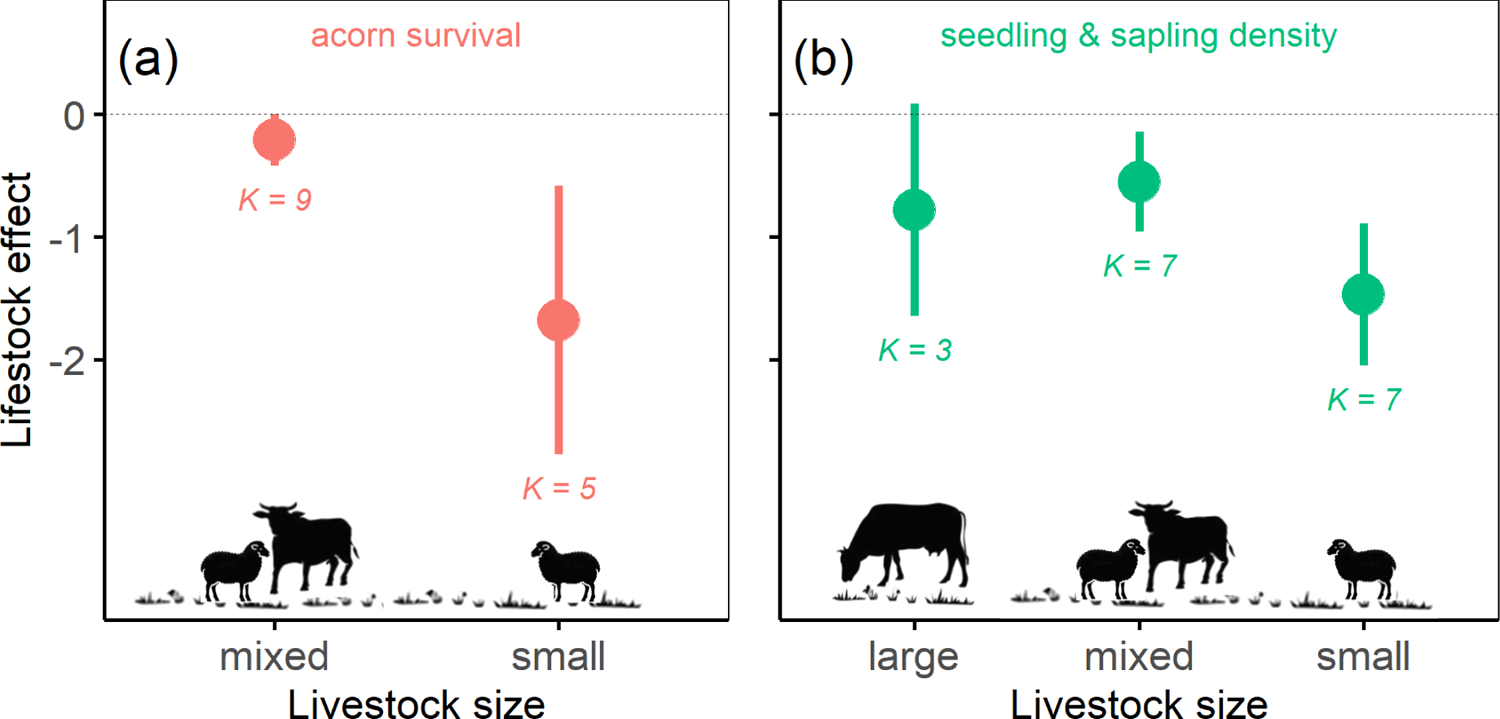
Mean effect size of domestic livestock, categorized by body size, on acorn survival (a) and on seedling and sapling density (b). Refer to Figure 2 for more explanations. No study eligible for meta-analysis has analyzed the effect of large livestock on acorn survival.

#### 3.3.2. Effects on oak seedling and sapling growth

Among the 13 case studies examining the impact of livestock on oak seedling and sapling growth, a very high negative effect was observed (−1.36; [−1.79, −0.94]). Importantly, this effect was not influenced by the early life stage of the oaks (*Q*M = 0.002, *P* = 0.964) or the size of the livestock (*Q*M = 0.370, *P* = 0.831).

#### 3.3.3. Effects on oak seedling and sapling density

The 17 case studies assessing livestock’s impact on early oak density consistently revealed a negative effect (−1.01; [−1.41, −0.62]). This effect was not deemed related to the oak’s life stage (*Q*M = 0.023, *P* = 0.879) but was dependent on livestock size (*Q*M = 8.038, *P* = 0.018). Case studies involving small-sized livestock exhibited the most pronounced negative effect on plant density (−1.47; [−2.05, −0.89]), significantly differing from those involving mixed-sized livestock (*Z* = −2.8, *P* = 0.006) (Fig. 4; Fig. B.5, Appendix B).

## 4. Discussion

Our meta-analysis pioneers the assessment of domestic livestock impacts on early oak life stages in global *Quercus*-dominated agroforestry systems. This is the first meta-analysis to differentiate the effects of livestock body size on the survival of acorns, seedlings, and saplings, as well as the density and growth of oak seedlings and saplings. Drawing upon a synthesis of 54 case studies conducted in diverse regions across Europe, America, Asia, and North Africa, our findings reveal an expected negative effect of livestock on oak regeneration across all life stages. However, we observed heterogeneity in the effect sizes among the studies, suggesting that factors beyond merely the presence or absence of livestock likely contribute to these variations. The overall effect size, quantified using Hedge’s *d*, substantiates the significance of the endeavor, with an effect size of −0.87 (95% CI: [−1.12, −0.62]). Our results contribute to a better understanding of the oak regeneration issues involved in silvopastoral systems across the globe.

### 4.1. Effects on early oak life stages

Our synthesis reveals that livestock affects negatively oak regeneration at the early developmental stages, encompassing acorns, seedlings, and saplings. For acorns, this is consistent with Leal et al. (2022), who emphasized the greater impact of wild ungulates on their survival compared to domestic livestock in Mediterranean woodlands. Our study did not examine interactions with wild ungulates or other acorn predators such as rodents and birds, which may have contributed to the observed variability in the effect on acorns across the case studies. While wild ungulates, including wild boar (Gómez and Hódar, 2008; Arosa et al., 2015) and deer (Weckerly, 2004), generally consume more acorns than domestic livestock, apart from pigs, the latter also partake in acorn consumption (Leiva and Fernández-Alés, 2003; Froutan et al., 2015; Mekki et al., 2019). Furthermore, the presence of cattle also influences rodent-mediated acorn predation and dispersal patterns (Muñoz and Bonal, 2007; Vaz et al., 2024).

The overall impact of livestock on oak seedlings, though similar in mean effect size to that on acorns, exhibited great variability across 29 case studies. This included four case studies where livestock had a positive effect. Although most studies indicated higher seedling survival in grazing-excluded areas compared to grazed areas, grazing may generate new ecological niches and enhance seedling survival (e.g., Etchebarne et al., 2016; Moradi et al., 2021), which may explain some positive effects. Also, enhanced seedling survival due to reduced competition from grasses, as a result of livestock grazing, can be an indirect positive effect of livestock presence. Additionally, oak seedlings may resprout in response to livestock browsing (Zhang et al., 2019; Vaz et al., 2019), though their sprouting ability can vary with livestock management practices and stocking rates (Pulido et al., 2010), oak species (Fortuny et al., 2020), and local environmental conditions (Jones, 2000; Plieninger et al., 2004). Broadly, the effects of livestock depend on the recruitment niche of each oak species, which is generally conditioned by the climate and the local environment. In Mediterranean oaks, recruitment tends to be promoted under shrub cover, which helps to overcome severe summer droughts. In agroforestry ecosystems with deciduous oaks (e.g., *Quercus robur*), the clearing of shrubs by livestock can create a suitable regeneration site for light-demanding seedlings, for which there is less risk of desiccation during the summer.

Our synthesis uniquely distinguishes livestock impacts on oak seedlings and saplings, unveiling a ‘very high’ negative impact on saplings (Cohen, 1988). Livestock foraging and trampling not only lead to reduced sapling height but also compact the soil and deplete moisture and organic matter, adversely impacting their establishment (Laskurain et al., 2013; Moradi et al., 2021). Yet, seedling and sapling establishment may also be facilitated by nurse shrubs, namely legume species (Gómez-Aparicio et al., 2004). In such cases, if browsers feed preferentially on nurse shrubs, oak regeneration may be negatively affected. On the other hand, if shrub cover is predominantly competitive, then clearing or even browsing of competitive shrubs may contribute to regeneration (Caldeira et al., 2014). Additionally, while early tree growth benefits from the facilitation of nurse plants and microclimatic amelioration, recurring shrub-clearing for grazing creates an ever-changing microhabitat that hinders sapling growth (Gómez-Aparicio et al., 2008), and saplings with limited defenses may become preferred forage for livestock (Göldel et al., 2016).

### 4.2. Impact of livestock body size and foraging behaviors

Our 54 case studies reveal that livestock body size, ranging from smaller animals like sheep to larger ones like cattle, can significantly affect the early oak life stages in agroforestry systems. Studies indicate that small-sized livestock, particularly sheep and to a lesser extent goats (notably, only two case studies exclusively involve goats), have more detrimental effects on oak regeneration, from acorns to saplings, compared to those focusing on large-sized livestock. Yet, no studies specifically addressed acorn predation by large livestock. Conversely, our results reveal lower negative effects on oak regeneration in mixed grazing systems. Notably, these trends are stark in acorn survival, with smaller ruminants appearing more acorn-predatory compared to mixed grazing systems. Sheep compromise seedlings by browsing and, under high grazing pressures, by trampling, which leads to bare soil as they remove litter or moss (Laskurain et al., 2013; Rossetti and Bagella, 2014). Seasonal grazing variations, notably in Mediterranean climates, further modulate these impacts, with small ruminants often targeting oak seedlings and saplings when grass is scarce (Ferreira et al., 2013). Contrarily, cattle and equines are typically managed to graze during periods of abundant grass, such as spring (Menard et al., 2002; Celaya et al., 2007). This distinction aligns with our data, where studies on small-sized livestock indicated more pronounced negative effects on oak seedling and sapling densities than those on mixed-size livestock. However, factors like soil compaction, potentially more severe with cattle than sheep or goats, were not analyzed in our synthesis (Lai and Kumar, 2020).

Our meta-analysis shows that systems with mixed-size livestock in oak agroforestry systems are associated with reduced negative impacts on early oak stages, compared to systems with only small-sized or, to a lesser extent, large livestock. Previous related research has also shown that mixed-size livestock more effectively control shrub encroachment (e.g., Ferreira et al., 2013). In Sudan and Oman, small livestock like sheep and goats were found to be more harmful to young trees than mixed-size groups and larger livestock (Ball and Tzanopoulos, 2020; Mohammed et al., 2021). Mixed-size livestock also enhance biodiversity by promoting vegetation structural complexity, benefiting a wide variety of herbaceous and arthropod species (García et al., 2013). Moreover, maintaining a variety of animal sizes and foraging behaviors also offers greater economic flexibility to farms (Anderson et al., 2012).

The lessened impact in mixed-size livestock systems may be attributed to the specific animal combinations in our data. Nearly 70% of the mixed-size case studies exclusively involved cattle and sheep. Although both are grazers, their distinct foraging behaviors, stemming from eco-physiological adaptations, can create greater vegetation heterogeneity (Adams, 1975; Hodgson et al., 1991; Benavides et al., 2009), potentially influencing oak establishment less than in areas with only sheep. While cattle, with their mobile tongues and broad, flat muzzles, are less efficient in grazing short swards, sheep, possessing smaller mouths and longer, narrower muzzles, can take smaller, more selective bites (Hofmann, 1989; Vallentine, 2001; Gordon and Benvenutti, 2006). Yet, both cattle and sheep graze on the delicate leaves of oak seedlings and saplings. The remaining case studies within this mixed-size subgroup were split between combinations of cattle, sheep, and goats, and those involving horses, sheep, and goats, suggesting that the more intense impact of goats on acorn consumption and browsing of seedlings and saplings is moderated in these mixed-size arrangements. As for horses, although they are not ruminants like cattle, both species have a significant degree of grazing overlap (Ferreira et al., 2013), with similar grazing selectivity, although horses, with teeth pointing slightly forward, can graze closer to the ground. Horses and cattle also generally prefer flatter terrain compared to small ruminants (Catorci et al., 2012), making acorns and young oaks on slopes less vulnerable to these larger animals.

### 4.3. Implications for oak conservation and management

Our meta-analysis reveals that livestock generally have a negative impact on oak regeneration, though effects vary by body size and are less pronounced in mixed-body-size grazing systems. Our findings challenge the view by a few authors that replacing cattle with sheep is less harmful to young plants (e.g., López-Sánchez et al., 2016), revealing a more complex scenario. This complexity may partly stem from the underestimation of small livestock’s effects on acorns and the lack of specific studies on large livestock’s acorn predation. Our synthesis highlights the significant negative effect of small livestock, particularly sheep and goats, on acorn survival when compared to mixed grazing systems. To enhance oak regeneration, adopting measures such as the individual protection of saplings and creating fenced areas during acorn shedding, germination, and early growth seasons is crucial for reducing acorn over-predation and promoting survival and early growth (Vaz et al., 2019; Löf et al., 2021). While these strategies may reflect common management practices in some well-managed forest systems, our results underscore the importance of their effective application. Yet, any exclusion activities should balance with the essential role of livestock as a source of income in these systems (Moradi et al., 2021). On the other hand, attributing larger livestock size with greater impact may arise from conflating size with related management factors. Practices often linked with larger cattle, such as mechanical mobilization for larger grazing areas, more pronounced trampling, and potentially more impactful overgrazing, could be contributing to perceived size-related effects (Pinto-Correia and Azeda, 2017).

A mixture of grazers of various sizes, such as sheep and cattle, during peak grass availability can be an effective compromise for reduced competition with oak seedlings. However, applying this approach globally requires careful consideration due to limited data from regions like Asia and northern Africa. In some Euro-Mediterranean regions, a shift to mono-specific herds favoring cattle (Catorci et al., 2012) contrasts with our results, which suggest benefits of mixed-body-size grazing systems. Conversely, management practices like rotational grazing (Plieninger et al., 2003) and stimulating livestock movement throughout the property (López-Sánchez et al., 2014), contribute significantly to sustainable oak regeneration. Traditional practices such as transhumance (Moreno and Pulido, 2009; Carmona et al., 2013) might not always fit modern contexts. Moreover, it is important to consider how livestock size and foraging behaviors interact with climatic conditions, including drought, for oak recruitment (Köchy et al., 2008). Stocking rates also influence this process and should be kept at low to intermediate levels to reduce negative impacts (López-Sánchez et al., 2014).

The heterogeneity detected in the effect sizes across studies examining the impacts of livestock on oak regeneration underscores the complexity of these effects. Crucially, livestock density plays a decisive role (see McEvoy et al., 2006), regardless of livestock type or size. For instance, a lower density of goats and sheep is likely less detrimental to oak regeneration compared to a very high density of cattle. Additionally, numerous factors such as breed type, livestock age, alternative forage, timing and duration of grazing, and the level of adaptation to young trees also significantly influence grazing impacts (Öllerer et al., 2019). These variables go beyond the simple presence or absence of livestock, which is a common element in all studies included in this meta-analysis, and they may contribute to the observed variability in effects.

### 4.4. Future research directions

Our analysis highlights imbalances in livestock impact studies eligible for meta-analysis on oak regeneration. Density research predominantly covers seedlings, with sapling studies being limited. Conversely, sapling growth is slightly more examined than seedling growth. Although acorn survival is relatively well-researched, the specific impacts of large livestock like cattle are overlooked. This indicates a need for broader research across oak life stages.

The scarcity of studies in certain regions corresponds to a gap in understanding the effects of varied domestic livestock. The limited focus on goats and the absence of other livestock in our synthesis are surprising, apart from cattle, horses, and sheep.

Furthermore, all studies addressing the effects of larger livestock have exclusively examined cattle. This highlights a need for broader, more inclusive research to encompass a wider range of domestic livestock and their impacts on oak agroforestry systems.

## 5. Conclusions

This meta-analysis evaluates livestock impacts on oak regeneration in *Quercus*-dominated agroforestry, distinguishing effects on acorns, seedlings, and saplings. Our findings show a high negative effect on early oak life stages linked to livestock size and foraging behaviors. Mixed-size systems mitigate these impacts better than single-size (especially small) livestock systems, as smaller livestock like sheep and goats particularly negatively impact acorn survival and seedling/sapling density. Our analysis challenges the idea that replacing cattle with sheep is less harmful to early oak stages, revealing the underappreciated impact of smaller livestock on acorn survival. Oak regeneration strategies should continue to employ established measures such as sapling protection in well-managed systems. Additionally, integrating livestock of diverse sizes and foraging behaviors, as well as establishing livestock-excluded areas during acorn seasons, can further enhance these practices and improve outcomes. Our synthesis urges the importance of expanded research in diverse regions and on varied livestock to refine agroforestry management globally.

## Authors’ contributions

**Abdullah Wadud:** Investigation, Formal analysis, Writing - Original Draft **Miguel Bugalho:** Writing - Review & Editing, Supervision **Pedro Vaz:** Conceptualization, Formal analysis, Writing - Original Draft, Writing - Review & Editing, Supervision.

## Acknowledgments

We acknowledge the researchers whose data supported this meta-analysis. AIW (PT/BD/143139/2018; SUSFOR—Sustainable Forests and Products Doctoral Program PD.00157.2012; CEABN-InBIO DOI 10.54499/UIDB/50027/2020; InBIO DOI 10.54499/LA/P/0048/2020) and PGV (SFRH/BPD/105632/2015; cE3c DOI 10.54499/UIDB/00329/2020; CHANGE DOI 10.54499/LA/P/0121/2020) were funded by FCT—Portuguese Science and Technology Foundation. We are deeply grateful for the valuable suggestions from three anonymous reviewers.

