## Supplementary data for "Meta-analysis of livestock effects on tree regeneration in oak agroforestry systems"

**Supplementary data for** Wadud AI, Bugalho MN, Branco M, Vaz PG. 2024. Meta-analysis of livestock effects on tree regeneration in oak agroforestry systems.

### Appendix A

#### Publication bias

Fail-safe numbers were consistently high across all meta-analyses. Using the Rosenthal approach, we determined that Fail-safe N greatly exceeded the threshold of 5 times the number of study cases plus 10 (Verdú et al., 2005). For instance, in the analysis of 24 case studies examining the combined impact of livestock on the survival of acorns, seedlings, and saplings, the Fail-safe N was 9272, surpassing the reference value of 130 (5 × 24 + 10 = 130) by 71 times. Additionally, funnel plots displayed no asymmetries, indicating the absence of publication biases in the underlying analyses.

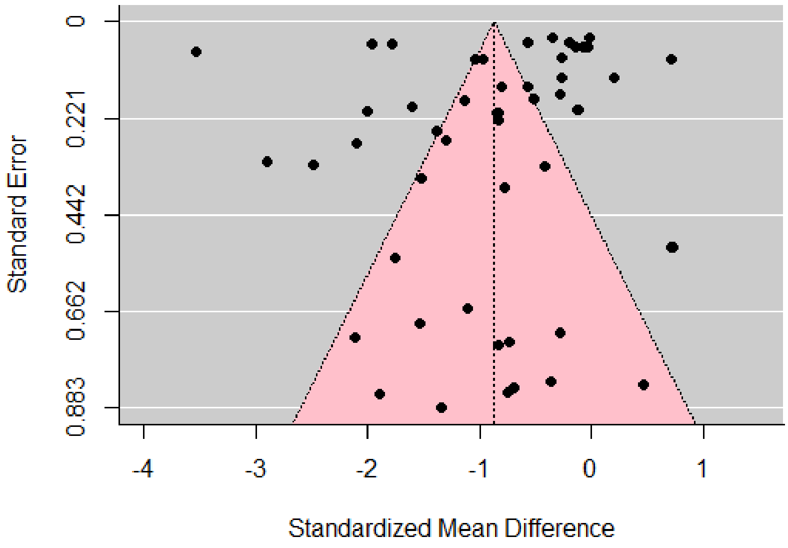


**Fig. A.1.** Funnel plot showing mean effect size versus precision, indicating asymmetry and possible publication bias in the study results.

### Appendix B

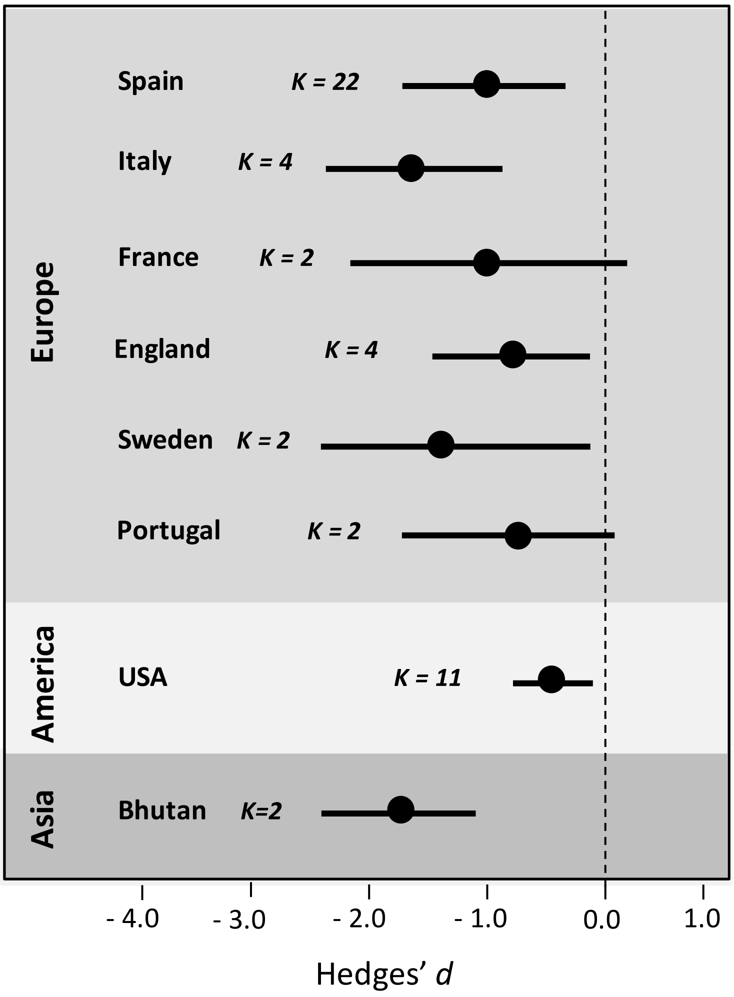


**Fig. B.1.** Forest plot illustrating livestock effect sizes and their 95% confidence intervals, categorized by continent and country. Canada, Greece, Algeria, Iran, and Israel have been omitted due to each having only one case study.

**Supplementary data for** Wadud AI, Bugalho MN, Branco M, Vaz PG. 2024. Meta-analysis of livestock effects on tree regeneration in oak agroforestry systems.

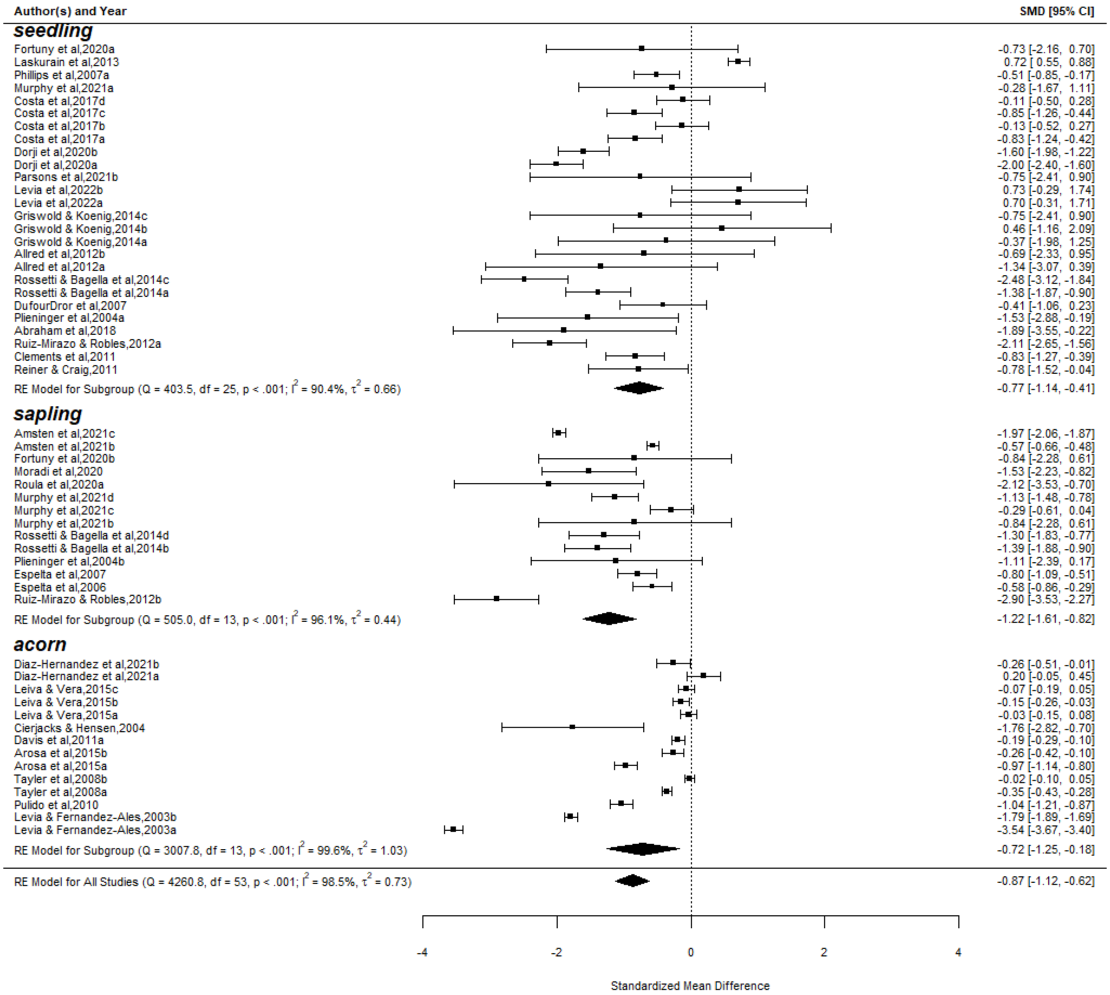


**Fig. B.2.** Forest plot illustrating livestock effect sizes and their 95% confidence intervals, grouped by early oak life stage.

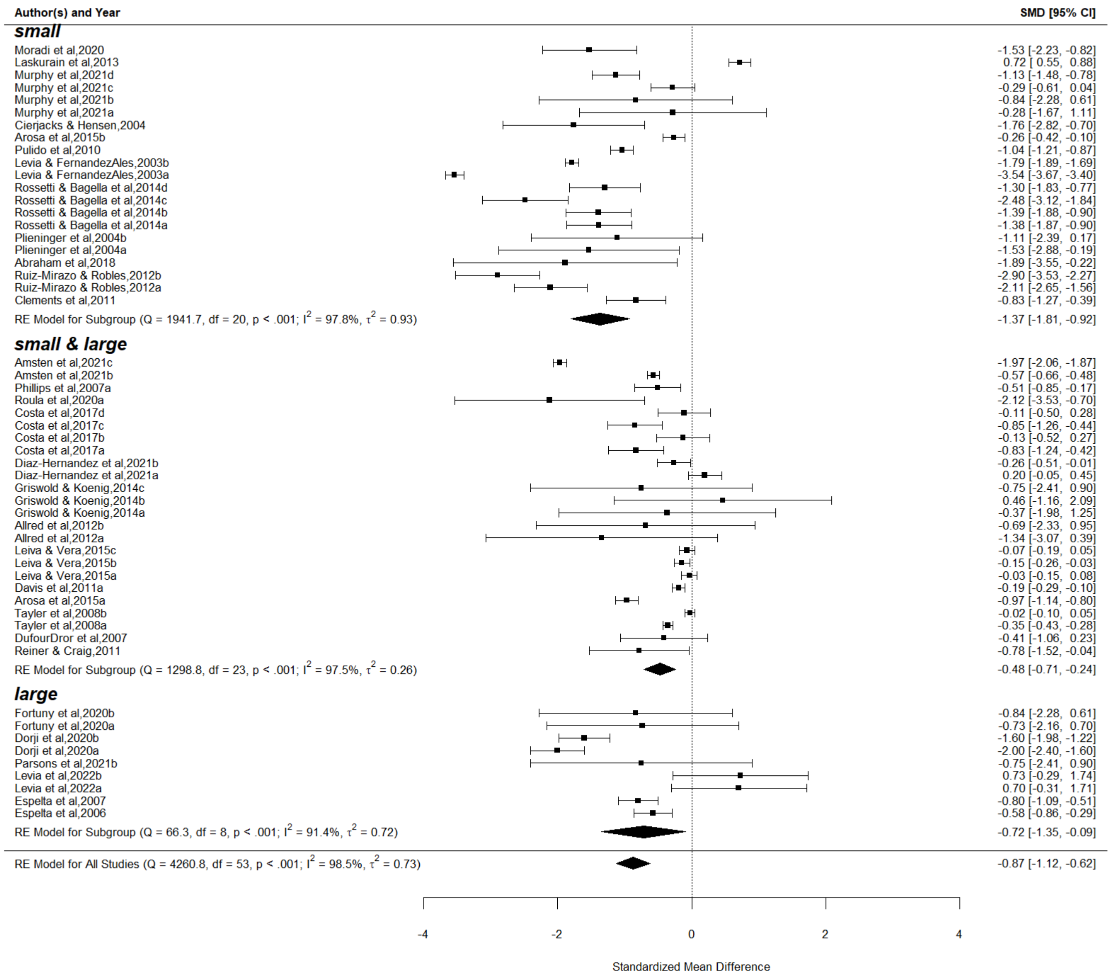


**Fig. B.3.** Forest plot illustrating livestock effect sizes and their 95% confidence intervals, grouped by livestock size.

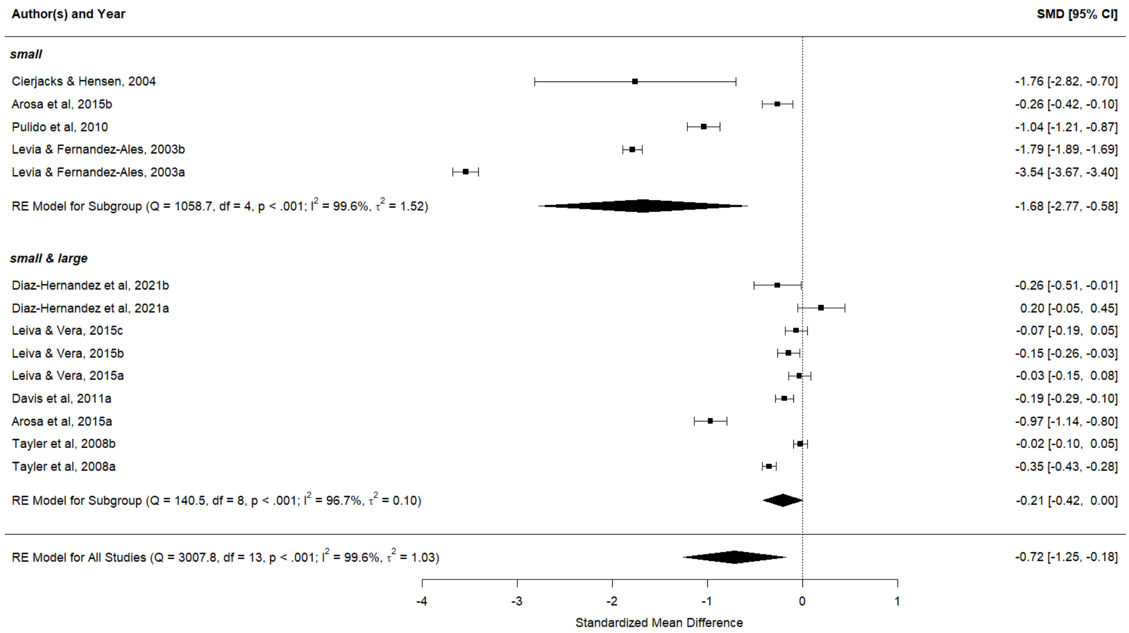


**Fig. B.4.** Forest plot illustrating livestock effect sizes on acorn survival and their 95% confidence intervals, grouped by livestock size.

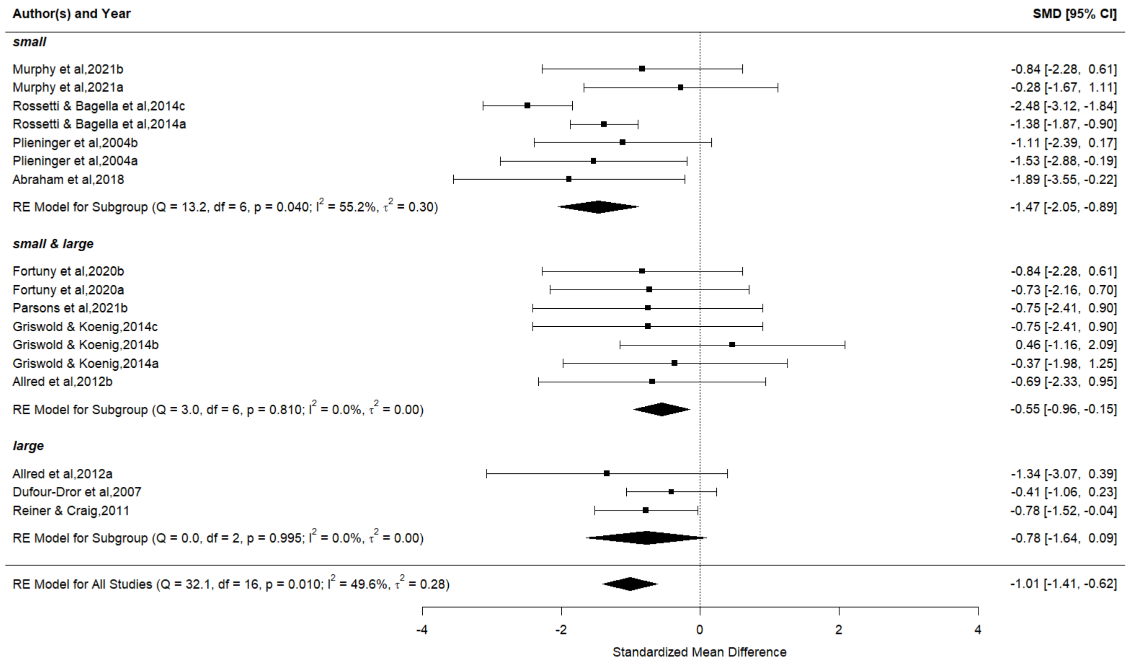


**Fig. B.5.** Forest plot illustrating livestock effect sizes on early oak density (seedlings and saplings) and their 95% confidence intervals, grouped by livestock size.
